# Reproductive success of Eastern Bluebirds (*Sialia sialis*) varies with the timing and severity of drought

**DOI:** 10.1101/575647

**Authors:** Reneé E. Carleton, John H. Graham, Adel Lee, Zachary P. Taylor, Jon F. Carleton

**Author notes:** These authors contributed equally to this work (Reneé E. Carleton, John H. Graham, Adel Lee). These authors also contributed significantly to this work (Zachary P. Taylor, Jon F. Carleton).

## Abstract

Drought affects avian communities in complex ways. We used our own and citizen science-generated reproductive data acquired through The Cornell Lab of Ornithology’s NestWatch Program, combined with drought and vegetation indices obtained from governmental agencies, to determine drought effects on Eastern Bluebird (*Sialia sialis* L.) reproduction across their North American breeding range for the years 2006–2013. Our results demonstrate that some aspects of bluebird reproductive success varies with the timing and severity of drought. Clutch size was unaffected by drought occurrence or severity during or within one and two months of clutch initiation, but hatching and fledging rates decreased as drought severity increased. Drought conditions occurring one month prior to the month during which eggs should have hatched, and two months prior to the month nestlings should have fledged also reduced the numbers of fledged offspring. We also demonstrate the value of datasets generated by citizen scientists in combination with climate data for examining biotic responses at large temporal and spatial scales.

## Introduction

Drought in varying degrees of severity and duration has affected the North American landscape and its biota for many hundreds of years [1, 2]. The lack of available soil moisture typical of drought can reduce or eliminate vegetation serving as either food or habitat for birds and their prey [3–5]. Consequently, it can negatively influence breeding birds dependent on those resources for their survival and reproduction. Unsurprisingly, breeding birds exhibit a variety of responses to drought, which may include a reduction in reproductive success [3, 6-8]. Dispersal away from drought stricken areas [9], increased mortality of adults, their offspring, or both adults and offspring, nest abandonment [3, 10], and reduced breeding attempts [3, 7, 10], have all been reported as a direct result of drought. Albright et al. [8], for example, found drought to have diverse effects on bird species abundance, with long-distance migratory species and those residing in semi-arid areas more severely affected than montane species, short-distance migrants, or synanthrophes.

Prediction models for the 21st century indicate drier conditions and periods of persistent drought for North America [11, 12]. Models also suggest that more than 50% of North American bird species will lose half of their distributional range, partly because of climate-associated impacts [13], such as drought. Moreover, because we know that drought negatively impacts many bird populations [7, 14, 15], it is critical to understand how and when drought will impact future reproduction across the entire breeding range of a species.

Detecting changes in species abundance at large spatial and temporal scales, such as the breeding range of a species across multiple years, requires intensive data collection. Few researchers, however, have the time or resources to accomplish such an undertaking. Citizen science, the collection of data by a network of volunteers, is increasingly used as a means of acquiring large data sets over wide geographic areas and long time periods [16–19]. Although some scrutiny of citizen science data is advisable, proper guidelines can provide for the generation of reliable data [20]. For example, the revelation of a climate-related change in the egg-laying dates of Tree Swallows (*Tachycineta bicolor*) was made possible by use of citizen science data [21]. One highly successful citizen science project is The Cornell Lab of Ornithology’s NestWatch Program, officially launched in 2008; it had evolved from the Cornell Nest Record Card program begun in 1965 [22]. NestWatch volunteers record breeding variables for 600 North American breeding bird species. These variables include number of nesting attempts, eggs produced, and young hatched and fledged, as well as nest location information obtained using online mapping applications [22]. Since the inception of NestWatch, more than 60 peer-reviewed articles using data generated by citizen nest observers have been published [23]. Large datasets generated by citizen scientists, in conjunction with standardized climate indicators, are therefore ideally suited to examine climate impacts across the range of a species.

The aim of this study was to use citizen science-generated data from NestWatch, and our own data, in conjunction with two drought indicators, the normalized difference vegetation index (NDVI) and North American Drought Monitor (NADM) PDSI-based drought levels, to determine how and when droughts of varying severity affect reproduction across the Eastern Bluebird (*Sialia sialis* L.) breeding range. As secondary cavity-nesting birds, Eastern Bluebirds readily adopt nest boxes provided them by individuals who enjoy hosting birds [24, 25] and, in some areas, nest boxes have replaced scarce natural nesting resources [25]. Inclusion of Eastern Bluebirds as a species monitored under NestWatch provided us with a significant data source for this study.

Geographical variations of clutch size are typical of passerines [20, 26, 27] and especially multi-brooded species [28, 29]. We hypothesized that reproduction of Eastern Bluebirds should follow this pattern but, drought during critical pre-breeding and breeding periods would negatively impact bluebird reproductive success. Our results indicated that drought conditions, regardless of severity, and occurring either pre-breeding or during the clutching period, had no significant effect on clutch size, but the hatching and fledging rates decreased as severity of drought increased. We also found that drought conditions occurring one month prior to hatching and two months prior to fledging also had a negative effect on Eastern Bluebird reproductive success.

## Methods

### Reproductive data

We obtained NestWatch (https://nestwatch.org/explore-data/) observation records for Eastern Bluebirds across their breeding range for the years 2006 through 2013. Records included a unique nesting attempt identification number, observer identification number, US state or Canadian province location, latitude and longitude of the nest under observation, year of observation, date of observation, clutch initiation date, clutch size, number hatched, number of young fledged, fledge date, and whether the nesting attempt was successful or unsuccessful. Following importation of all records into a Sequel Query Language (SQL) database, we ran queries to identify and eliminate records of nesting attempts and associated activity outside of the typical March through August breeding season, records where more than 7 eggs were observed in a single nest that might indicate use by more than one female or reuse of a nest containing an abandoned partial clutch, and records where the number of eggs was recorded as 0 but numbers of young, or young fledged was recorded as greater than 0. We also eliminated records for which the recorded fledge date was greater than 40 days past the clutch initiation date; given an expected sequential oviposition, an average 12 to 15 days of incubation, and 20 days between hatch to fledge [30], we judged the reliability of these observations as questionable. Records from states on the breeding range boundaries with fewer than 5 submissions were also deleted as well as those with geographic locations outside the range of Eastern Bluebirds or otherwise questionable. We included data of the same NestWatch reproductive variables and time period from our study site located on our home institution’s land tract in Floyd County, GA, where we have monitored Eastern Bluebird reproduction since 2002.

#### Ethics Statement

Our data collections were carried out in strict accordance with the Ornithological Council’s recommendations [31] and under the permits issued by the US Geological Survey’s Bird Banding Laboratory and the State of Georgia’s Department of Natural Resources. The protocol was approved by the Institutional Animal Care and Use Committee of Berry College (Protocol Number: 2006-13-005).

### Drought indices

#### NADM drought categories

We used latitude and longitude recorded for each nest box to identify the county in which it was located and for the nearest weather station within 40 km. We then downloaded archival drought status categories by county and for each month during the years 2006–2013 from the NADM website (http://drought.gov; see also for an explanation of drought status and Palmer Drought Severity Index). For our analyses, drought status categories were treated as ordinal variables (Table 1).

**Table 1.** Drought categories, Palmer Drought Severity Indices (PDSI), descriptions, and possible impacts. Data source: National Drought Mitigation Center. 2016. Accessed November 2018. http://droughtmonitor.unl.edu/MapsAndData.aspx.

| Category | PDSI | Description | Possible Impacts |
| --- | --- | --- | --- |
| N | – | No drought |  |
| D0 | -1.0 to -1.9 | Abnormally dry | Going into drought: short term dryness slowing planting or growth of crops or pasture. Coming out of drought: some lingering water deficits; pastures or crops not fully recovered |
| D1 | -2.0 to -2.9 | Moderate drought | Some damage to crops or pastures; streams, reservoirs, or wells low |
| D2 | -3.0 to -3.9 | Severe drought | Crop or pasture losses likely; water shortages common; water restrictions imposed. |
| D3 | -4.0 to -4.9 | Extreme drought | Major crop/pasture losses; widespread water shortages or restrictions |
| D4 | -5.0 and less | Exceptional drought | Exceptional and widespread crop/pasture losses; shortages of water in reservoirs, streams, and wells, creating water emergencies |

#### Normalized Difference Vegetation Index

NDVI (normalized difference vegetation index) is correlated with net primary productivity (NPP) [32]. We calculated a NDVI value for each site for each month during the nesting season using 14-day Advanced Very High Resolution Radiometer (AVHRR) 1-km composites downloaded from the United States Geological Survey. For each month, we averaged two 14-day composites and determined the NDVI for each nesting site using ArcMap 10.3 (ESRI, Redlands, CA). Standardized NDVI was calculated, following Albright et al. [32], as 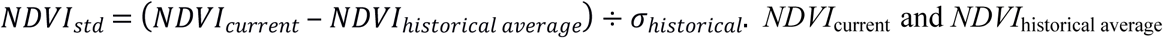 are for specific locations. *NDVI*_std_ makes it possible to compare widely different sites, such as relatively arid and relatively humid locales.

### Statistical analyses

The primary question we wanted to address was, “To what degree was Eastern Bluebird reproductive success impacted by drought?” We conducted 3 separate analyses (generalized linear mixed-effect models) in R’s lme4 package, with clutch size, hatch ratio (number of eggs hatched divided by clutch size), and fledge ratio (number of young fledged divided by clutch size) as respective response variables. To study the effect of drought on our respective outcome variables, we considered both current drought status and drought statuses shortly prior to the reproductive event of interest (hereafter referred to as “prior drought”). Current drought status was the drought status at the time of clutch initiation, hatching, or fledging of young, respectively, while prior drought status was that recorded at one month and two months prior to the date of clutch initiation, hatching, or fledging, respectively.

We set up our analysis as a tiered modeling approach. First, we specified a baseline null model (Model 0), which included the numerical month of clutching, hatching, or fledging (sequentially numbered from March through August), NADM climate region (Northeast, Southeast, South, and High Plains, with Midwest as the reference category; Table 2), latitude and longitude, and standardized NDVI. Our first-tier model (Model 1) added current drought status (D0 through D4, with N as the reference category) to the baseline explanatory variables. A second-tier model (Model 2) added both the current month and one-month prior drought statuses as explanatory variables. Our third-tier model (Model 3) included current month, one month, and two months prior drought statuses. We elected to only include drought conditions within two months of breeding as prevailing conditions 3 to 5 months prior to clutch initiation would occur during winter months when some populations had migrated to other regions or when breeding territories had yet to be established. We considered the drought status of one and two months prior to clutching, hatching, and fledging as possibly critical time periods for not only nest building, but also when insect populations should undergo increases which would in turn be important for the feeding of nestlings [33, 34].

**Table 2.** U.S. states and adjacent Canadian provinces included in North American Drought Monitor climate regions which fall within the Eastern Bluebird breeding range.

| Climate region | U.S. state and Canadian provinces |
| --- | --- |
| Northeast | CT, DE, MA, MD, ME, NH, NJ, NY, PA, RI, VT, WV, QB, ON |
| Midwest | IA, IN, IL, KY, MI, MN, MO, OH, WI |
| High Plains | KS, ND, NE, SD, MB |
| Southeast | AL, FL, GA, NC, SC, VA |
| South | AR, LA, MS, OK, TN, TX |

At each tier, we assessed whether the added drought status variable had a significant effect on the clutch size, hatching rate, and fledging rate. In particular, we did likelihood ratio tests to test the hypothesis that the drought status variable added at that tier had a non-zero effect. If the drought status variable had a significant effect, then we examined how the different levels of drought affected the reproductive outcomes. For this study, the focus was not to predict the outcome variable or to select the best subset of explanatory variables, but to instead test hypotheses about drought status variables, and to gain insights into the effect of different levels of drought severity and timing of drought (for significant drought status variables).

As described above, each of the baseline and three sequential models included a set of explanatory variables related to climate and certain NestWatch data categories. We centered the continuous variables month, latitude, longitude, and standardized NDVI around their respective means prior to model fitting. The NADM region in which nests were located served as a proxy for differences in seasonal breeding behavior across broad geographical areas. We included an interaction term between region and month in our models to capture differences as the breeding season progressed since reproductive attempts and success are known to vary within a season and across geographic areas [20, 28, 29].

In addition to the various fixed effects, we used random effects to account for sources of spatial variability not captured by the fixed effects. We included county in which nests were located and unique NestWatch user identification numbers nested within county as random effects. These random effects accounted for random variability on a larger spatial scale (i.e., county level) and the variability in skill level among the individuals that reported data to NestWatch. Recognizing that these spatial random variability patterns varied by year, we nested the above random effects within year. This allowed the model to reflect the spatial variability that would not necessarily be the same from one year to the next.

We used a generalized linear mixed-effects model (glmer function in the lme4 package for R) to conduct the above and Nelder-Mead estimates of the model parameters. For the analysis with clutch size as the outcome variable, we assumed a Poisson probability distribution, with a log, X*β* = ln(*µ*), link function. In contrast to clutch size, the assumptions of a Poisson distribution were violated for both the number of young hatched and the number of young fledged, because of the large percentage of zero observations. Rather than using the number (or count) of eggs hatched (or young fledged) as the outcome variable, we used the success rate (number of eggs hatched and number of young fledged divided by clutch size). With hatching and fledging ratio as the outcome variable, we fitted a generalized linear mixed model (glmer from package lme4), assuming that the hatching and fledging ratios were generated from a Binomial probability distribution, with the logit, *Хβ* = *ln* (µ/(1 ‒ µ)), as the link function. The response variable was a ratio (ranging between 0 and 1). We assessed the collinearity among the explanatory variables of each model by calculating the generalized variance inflation factor (GVIF) [35], using the R package car. We used the degrees of freedom (*df*) adjusted measure, GVIF^(1/(2 · *df*))^, provided by the vif function in car. We found that this adjusted measure was less than two for all the sets of explanatory variables considered, which indicated that the explanatory variables were not highly correlated.

## Results

### Drought conditions

Abnormally dry (category D0) to drought conditions of varying severity (categories D1– D4) occurred within Eastern Bluebird nesting territories throughout the study period (Fig. 1). No exceptional drought (category D4) occurred within the NADM Northeast region for years 2006– 2013. Periods of exceptional drought occurred in the South region between years 2006–2009 and 2011–2013. In 2011, the state of Texas, one of the South region states, experienced the worst one-year drought since the inception of record keeping in 1895 when approximately 90% of its land mass was declared to be in a state of exceptional drought (D4) and over 31,000 wildfires burned an estimated 4 million acres [36]. The Southeast region also experienced periods of exceptional drought during 2006–2007 and 2012–2013, with nearly 40% and up to 10% of the land area affected, respectively. Although the Midwest region experienced periods of extreme drought (D3) during 2006–2009, exceptional drought occurred only during 2012, when less than 10% of the land area was affected. In the High Plains region, abnormally dry (D0) conditions affected up to 100% percent of the land area during the 7 years examined in this study. Extreme (D3) and exceptional drought (D4) occurred there during the latter half of 2012 and for nearly all of 2013.

**Fig 1.**
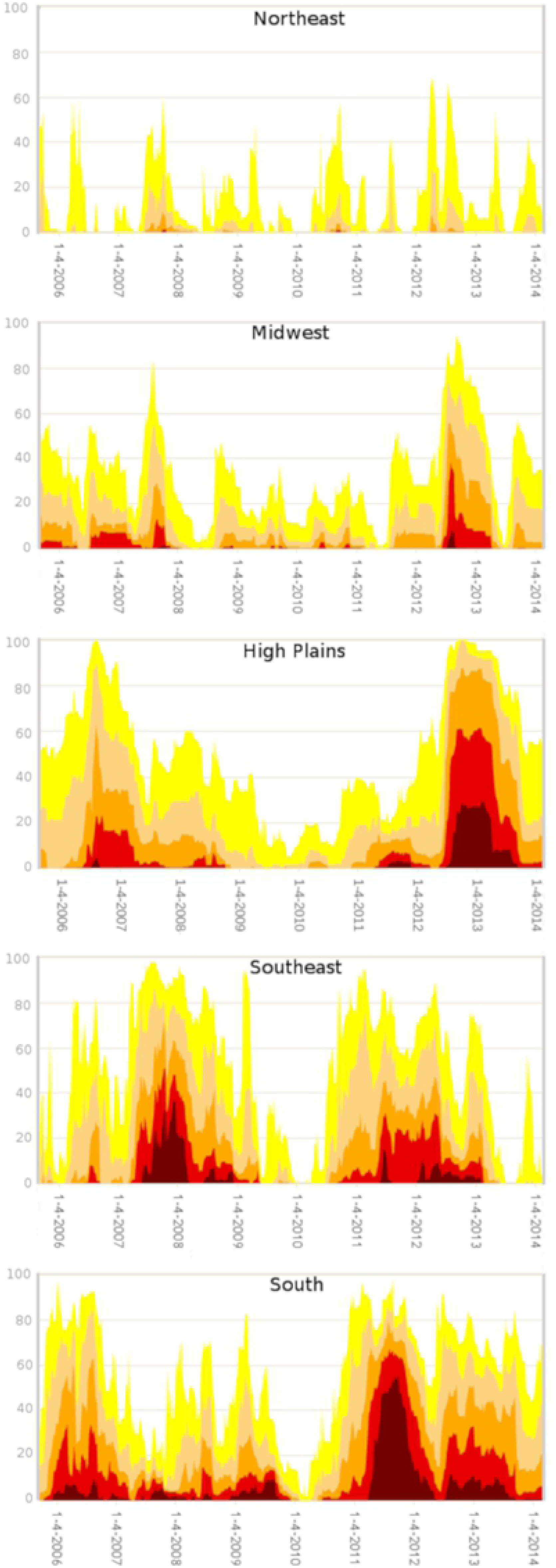
Percentage land coverage during drought conditions (2006─2013) by NADM climate regions encompassing the breeding range of Eastern Bluebirds (*Sialia sialis*). Modified from NADM climate region map archive (https://droughtmonitor.unl.edu/Maps/MapArchive.aspx).

#### Reproductive data

We received 24,368 individual nest observations records from NestWatch and included 708 observation records generated from our nest box monitoring program. Following elimination of records with inaccuracies, 21,574 records were used for the analyses (Fig. 2). The number of records was lowest in 2006 (*n* = 413), but increased yearly from *n* = 1876 in 2007 to *n =* 3,839 in 2012, before declining slightly in 2013 to *n* = 3,698.

**Fig. 2.**
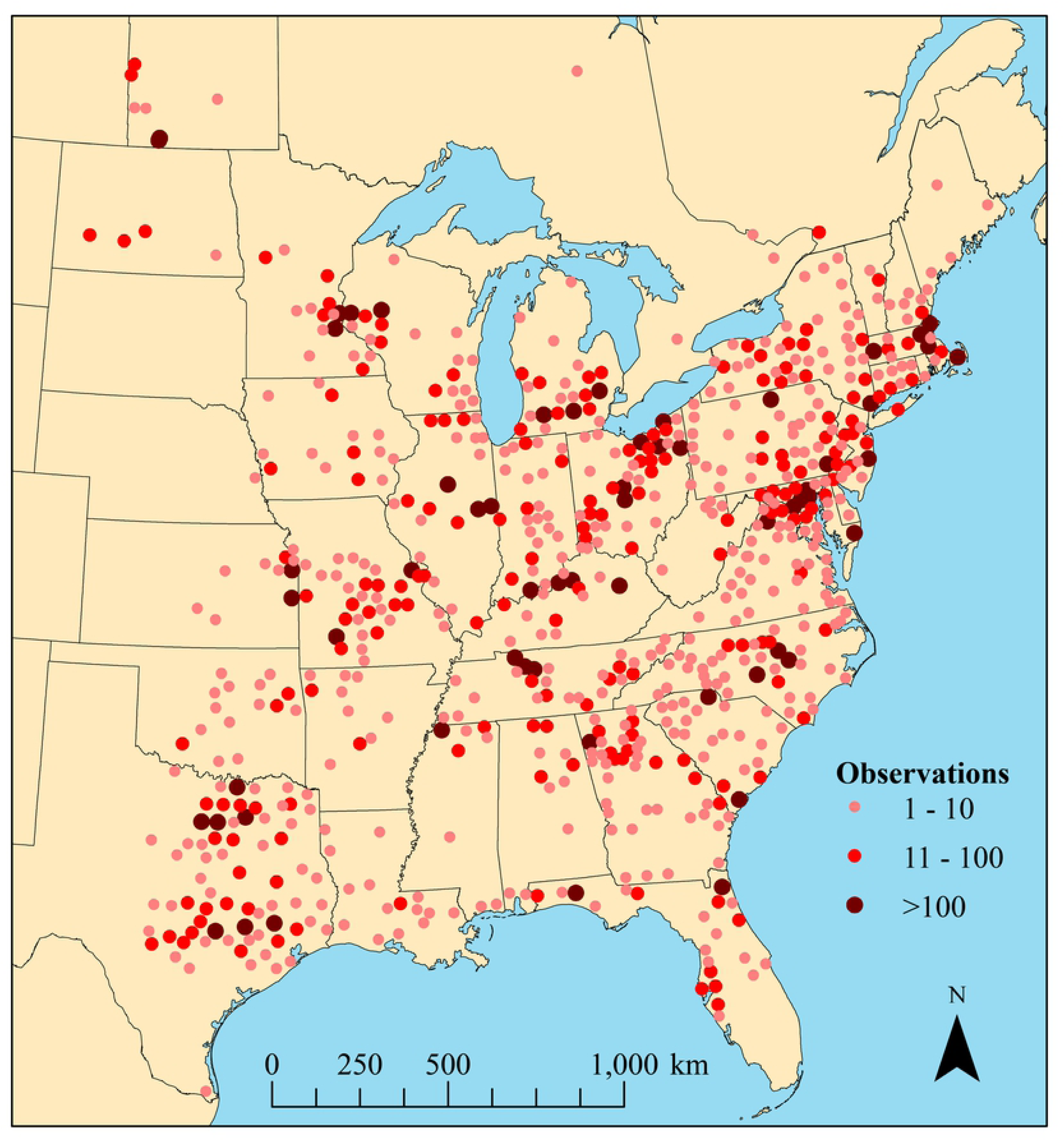
Observation sites within the North American breeding range of Eastern Bluebirds (*Sialia sialis*) aggregated by county (in US) or administrative region (in Canada), 2006– 2013.

#### Clutch size

Clutch size ranged from 1 to 7 eggs, with a mode of 5 and a mean of 4.37 (Fig. 3). To the left of the mode, the increase was nearly exponential. The drop-off above the mode of 5 was steep, with only a small fraction of the clutches having 6 eggs. The number of eggs per clutch did not decline with increasing drought severity (Fig. 4).

**Fig. 3.**
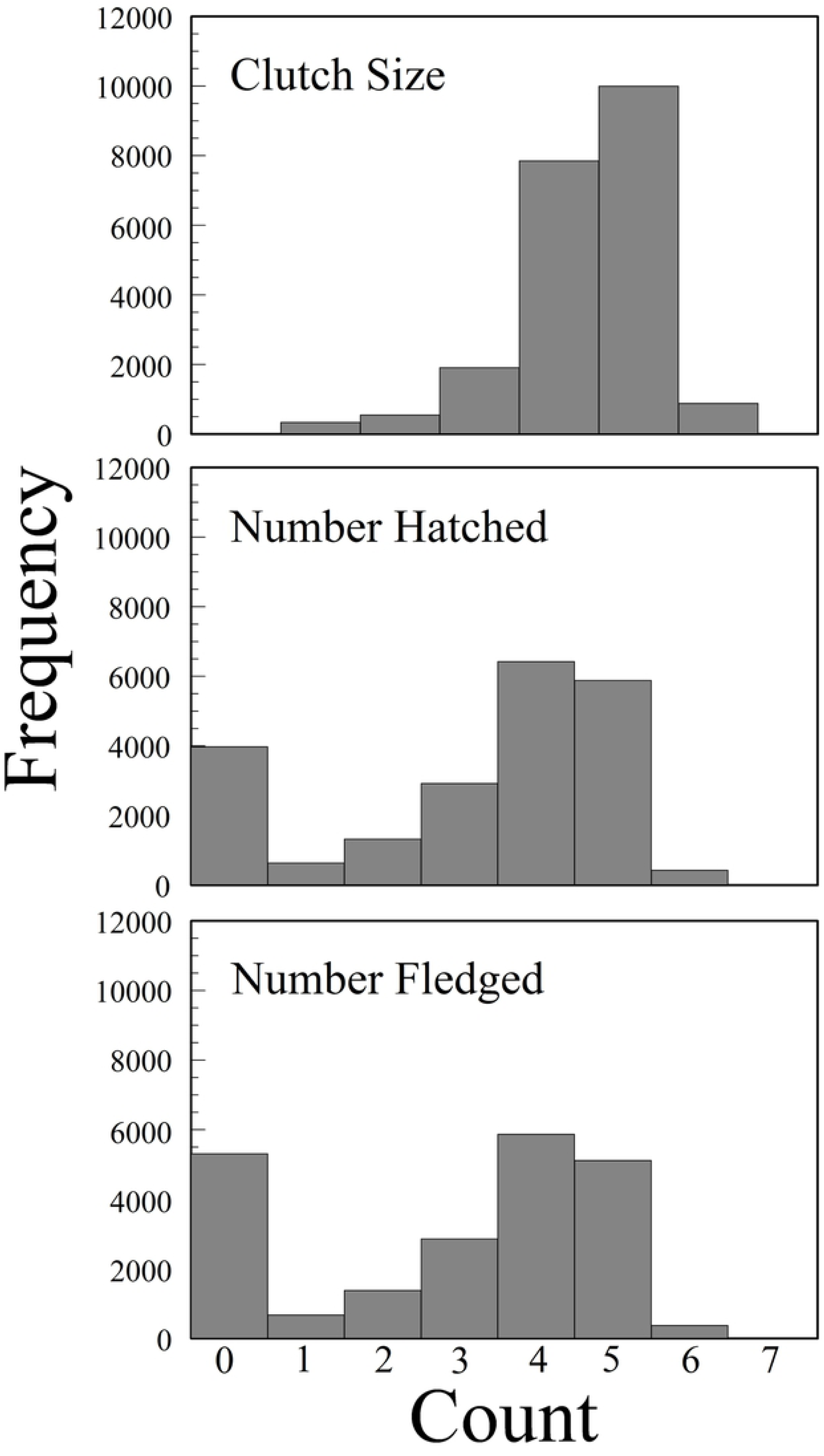
Frequency distributions of Eastern Bluebird clutch size, number hatched, and number fledged across their entire range in North America, 2006-2013.

**Fig. 4.**
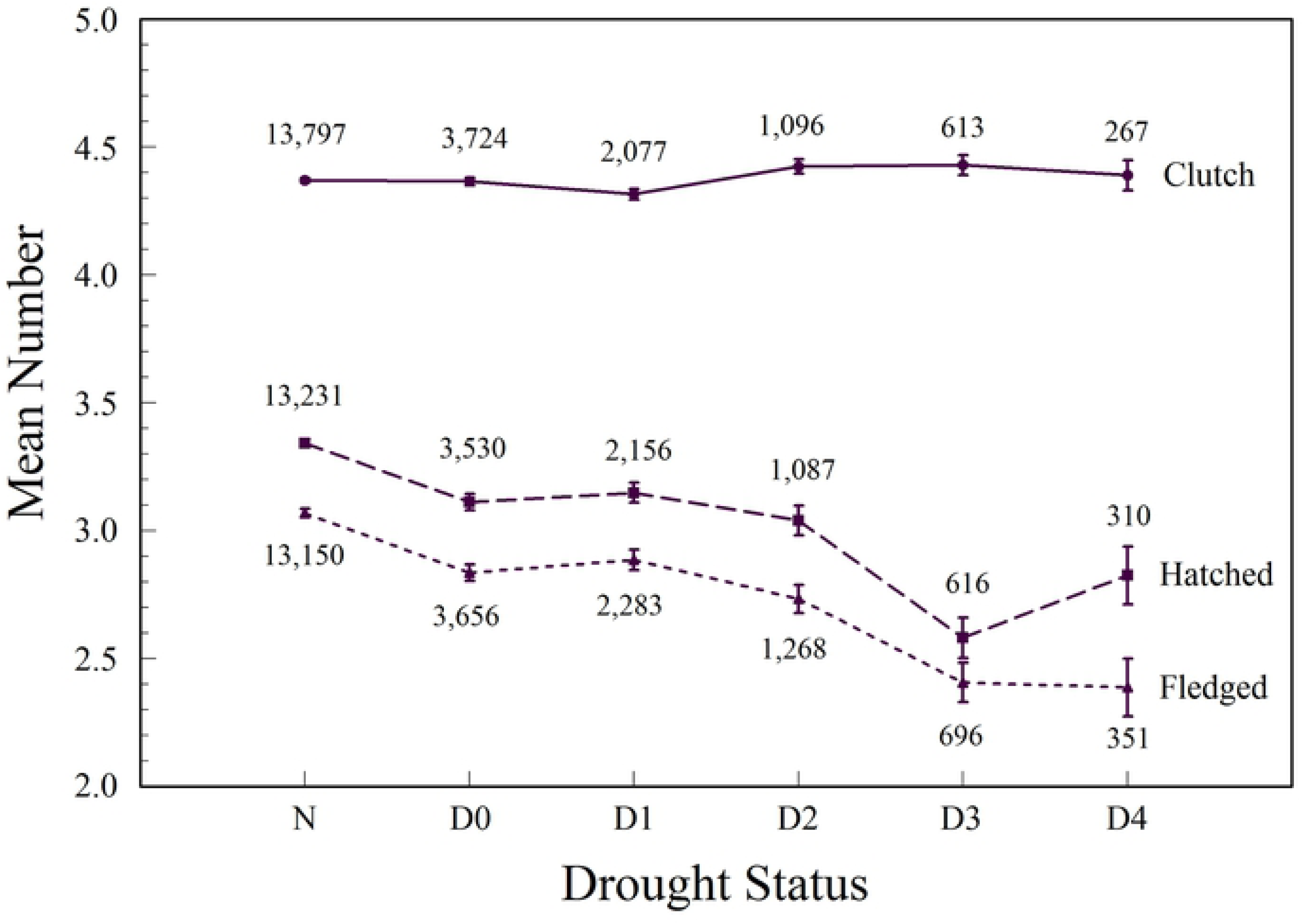
Mean number of Eastern Bluebird eggs laid (clutch size), eggs hatched, and young fledged by North American Drought Monitor drought status categories, 2006-2013.

We used likelihood ratio (LR) tests to compare the pairwise goodness of fit of the statistical models, a simpler null model (Model 0), and a series of alternative models (Models 1–3) with additional explanatory variables. The first LR test (Model 0 vs Model 1) tested whether the effect of the current month’s drought status was significantly different from zero (*H*_0_: *β* = 0). This test was executed by calling the R function anova, with the fitted Model 0 (null model with no drought status explanatory variables) and the fitted Model 1 (alternative model with drought status variables added to the base variables). There was no evidence to reject the null hypothesis (*Χ*^2^ = 1.30, *df* = 5, *P* > 0.90); the additional drought status variable did not improve the fit when added to Model 0. This result was clearly in line with the coefficient estimate results for drought statuses (D0–D4) during the months that the clutches were laid (Table 3). None of the drought status levels (D0–D4) had a significant effect on clutch size when compared with N (normal rainfall, no drought) (Fig. 4).

**Table 3.**
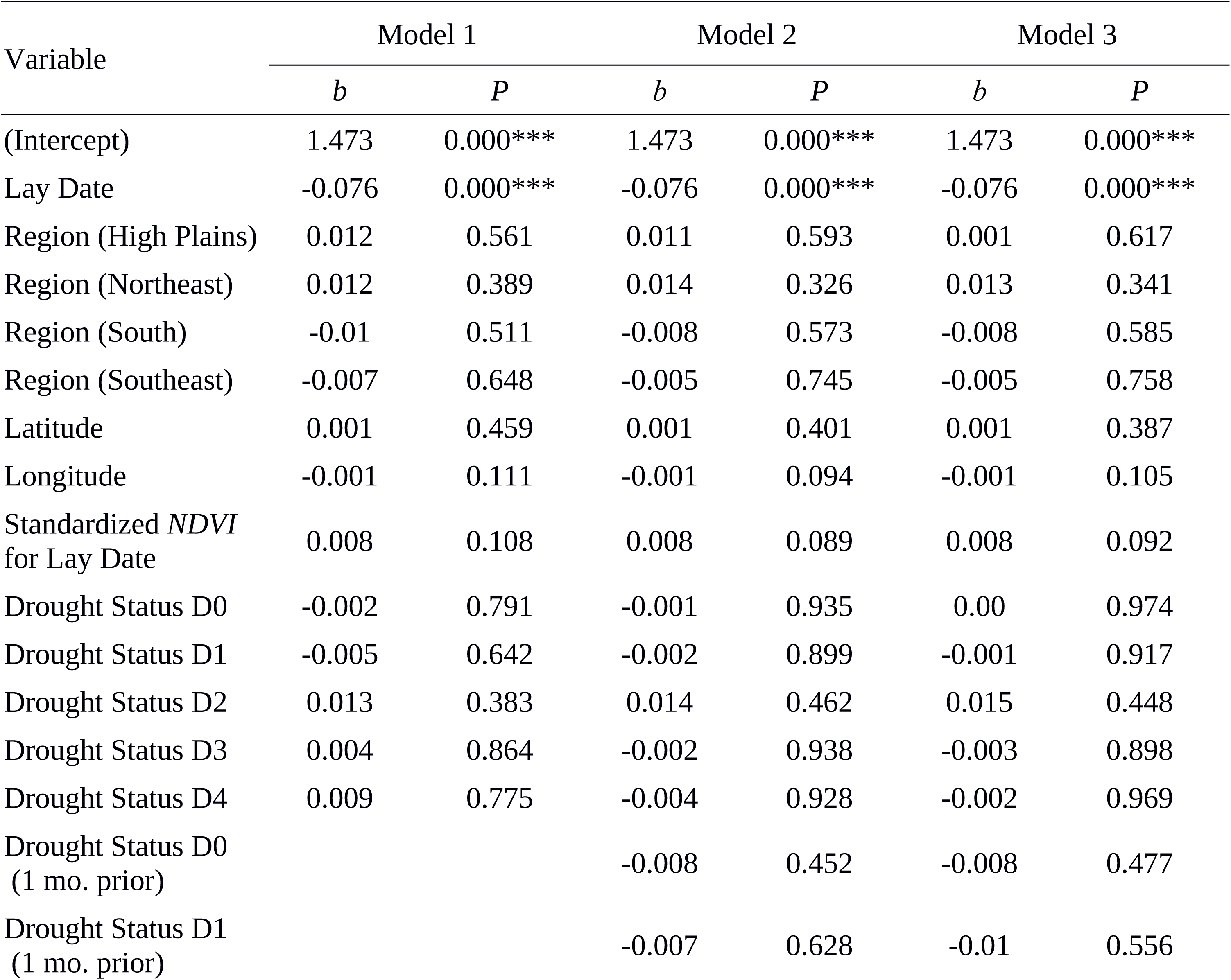
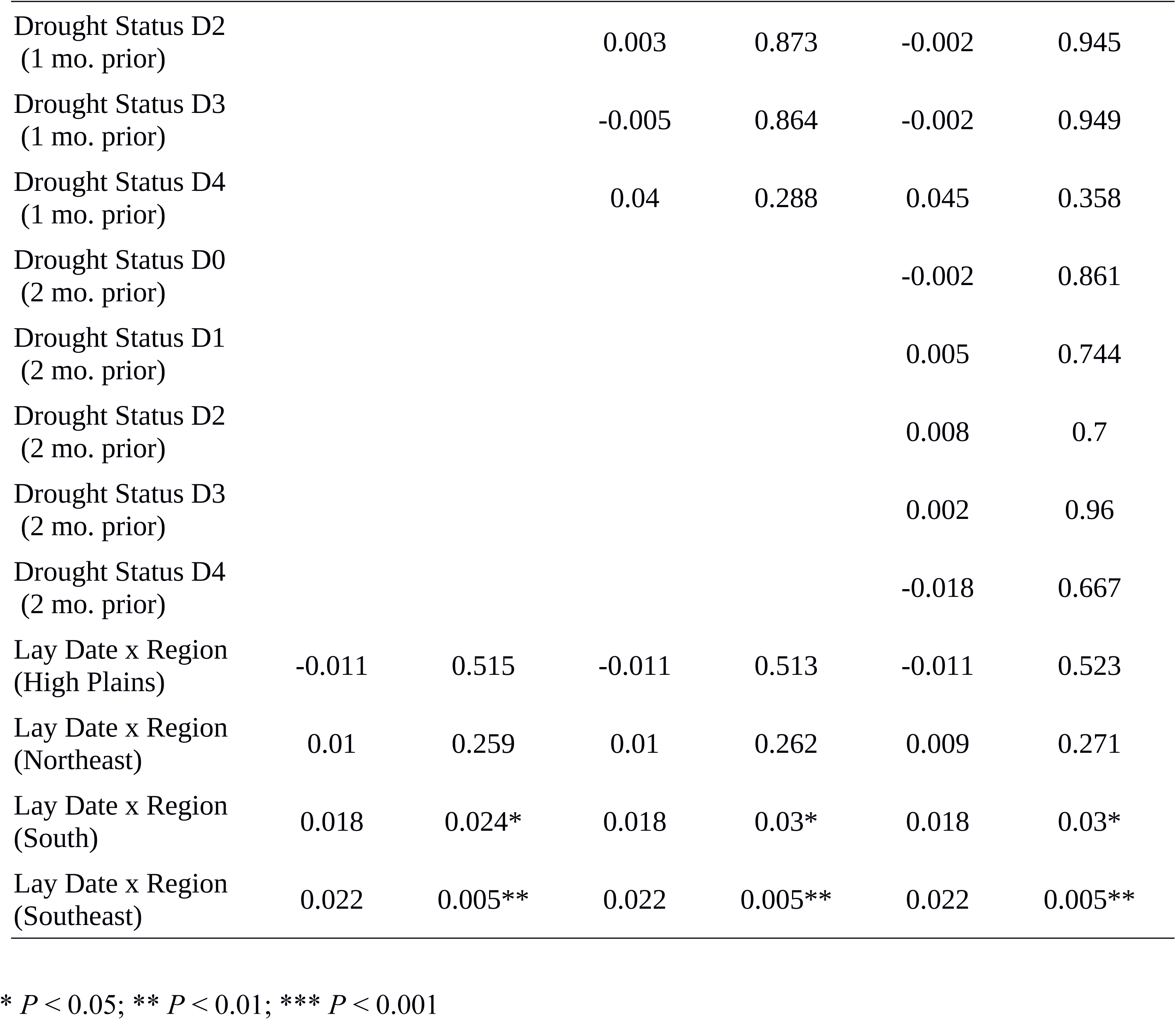
Parameter *β* estimates (*b*) and probabilities (*P*) for three generalized linear models used to evaluate drought effects on Eastern Bluebird clutch size. Explanatory variables are lay date, region, latitude, longitude, standardized *NDVI* for the lay date, drought status during the lay date, drought status one month prior to the lay date, and the interaction of lay date and region.

The second LR test, Model 1 vs Model 2, tested whether the effect of drought status one month before a clutch was laid was significantly different from zero (*H*_0_: *β* = 0), and the third LR test, Model 2 vs Model 3, tested whether the effect of drought status two months before a clutch was laid was significantly different from zero (*H*_0_: *β* = 0). Both tests had large *p*-values (*Χ*^2^ = 2.58, *df* = 5, *P* > 0.75 and *Χ*^2 =^ 0.66, *df* = 5, P > 0.95, respectively), indicating no evidence for the effect of prior drought status one or two months earlier on clutch size (Table 3).

We also tested whether a more complex random effects structure, with userid nested within county and county nested within year, provided a better fit than a simpler one with just year as a random effect. The LR test clearly favored the simpler random effects structure with just year as a random factor (*Χ*^2^ < 0.001, *df* = 2, *P* > 0.99). The results reported here (Table 3) were based on using the simpler structure with year as a random factor.

Exponentiated regression coefficients (*e^b^*) can help with the interpretation of the coefficients in a particular model. For example, using the parameter estimates for Model 1, which was not significantly different from Model 0, the effect of month (Lay Date) was significant, and the exponentiated coefficient (*e^b^*) for month, *e^b^* = *e* ^-0.076^ = 0.927, gives the multiplicative term used to estimate the clutch size when month increases by one unit (say April to May, or 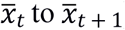). Thus, the multiplicative effect of month on clutch size, of a one-month increase, was 0.927. For example, if the hypothetical mean clutch size in April was 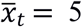 eggs, then the predicted clutch size for May would be 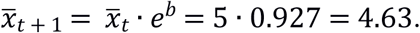 Clutch sizes peaked early in the breeding season and then declined as the season progressed (Fig. 5). There was a significant interaction between the South and Southeast regions and the month a clutch was laid. Compared to the Midwest region, there was an increasing multiplicative effect on clutch size for the South (*e* ^0.018^ = 1.018) and Southeast (*e* ^0.022^ = 1.022). Thus, if the mean clutch size for the Midwest was 5 in April, then in May it would be 5.091 for the South and 5.111 for the Southeast.

**Figure 5.**
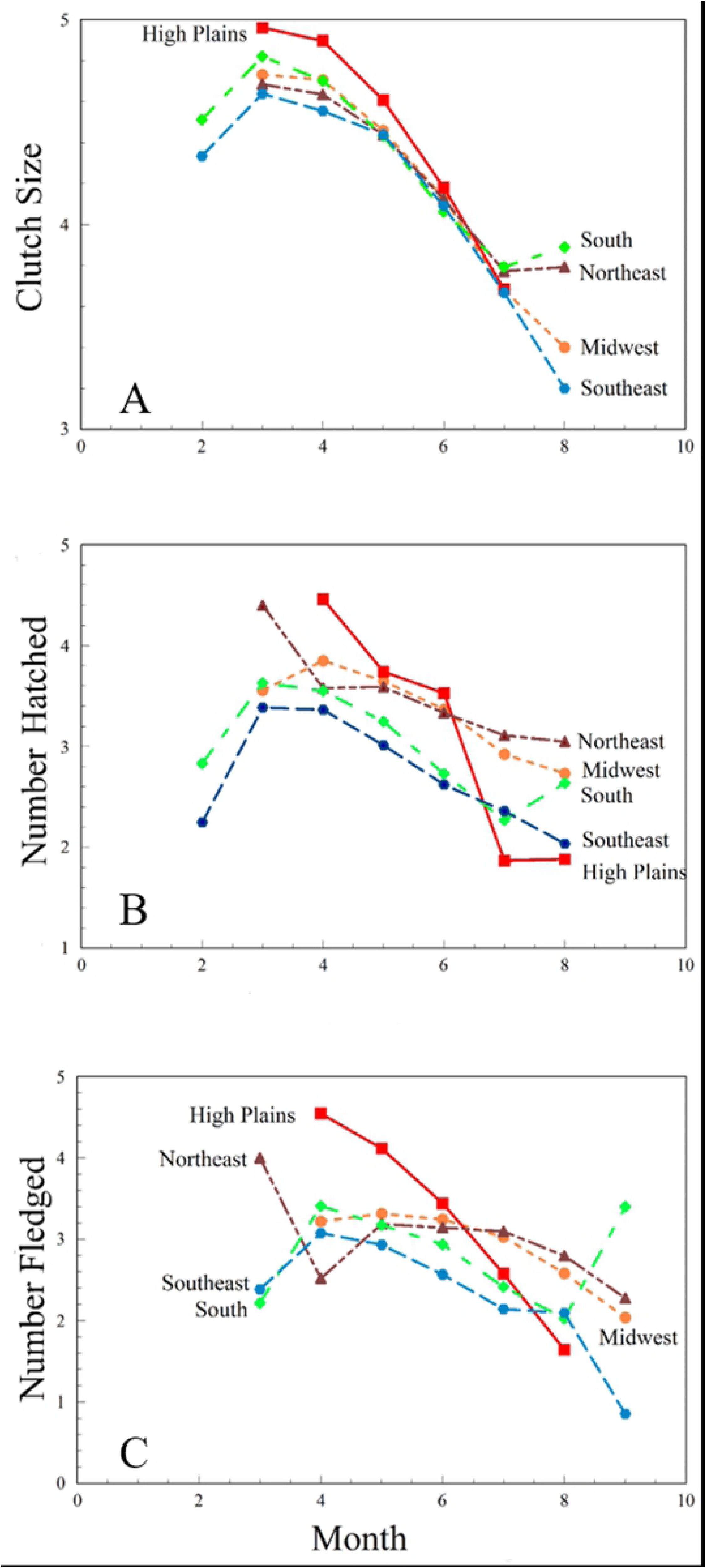
Variations in Eastern Bluebird clutch size, number of young hatched, and number of young fledged by North American Drought Monitor climate region and month, 2006-2013.

Because latitude and longitude are typically significant indicators of clutch size, we examined whether or not this trend was present in our data by removing the region variable from the null model. Latitude (*Z* = 2.66, *df* = 1, *P* < 0.01) and longitude (*Z* = −2.02, *df* = 1, *P* < 0.05), significantly affected clutch size when the region variable was not included.

#### Hatching rate

The modal number of hatching eggs was 4, followed by 5 and zero (Fig. 3). The distribution was bimodal. The average number hatching was 3.23. The number of hatched young declined from more than 3.3 to fewer than 3.0 hatching per nest with increasing drought status, especially for the two most extreme categories (Fig. 4).

As with clutch size, we also did nested, pairwise likelihood ratio tests comparing the models. Model 1 was a significant improvement over Model 0 (*Χ*^2^ = 33.00, *df* = 5, *P* << 0.0001). Likewise, Model 2 was a significant improvement over Model 1 (*Χ*^2^ = 45.2, *df* = 5, *P* << 0.0001), and Model 3 was a significant improvement over Model 2 (*Χ*^2^ = 19.7, *df* = 5, *P* < 0.0014). We again tested, as with clutch as the response variable, whether a simplified random effects structure was warranted. The nested random effects structure (with userid nested within county and county nested within year) provided the best fit (*Χ*^2^ = 6684.5, *df* = 2, *P* << 0.0001).

We report the model fitting results for hatch ratio for Models 1, 2, and 3 in Table 4. Here, we interpret the drought status results for Model 3, since our sequential hypothesis testing results were overwhelmingly in favor of including both the current and prior (one and two months) drought status variables in the model.

**Table 4.**
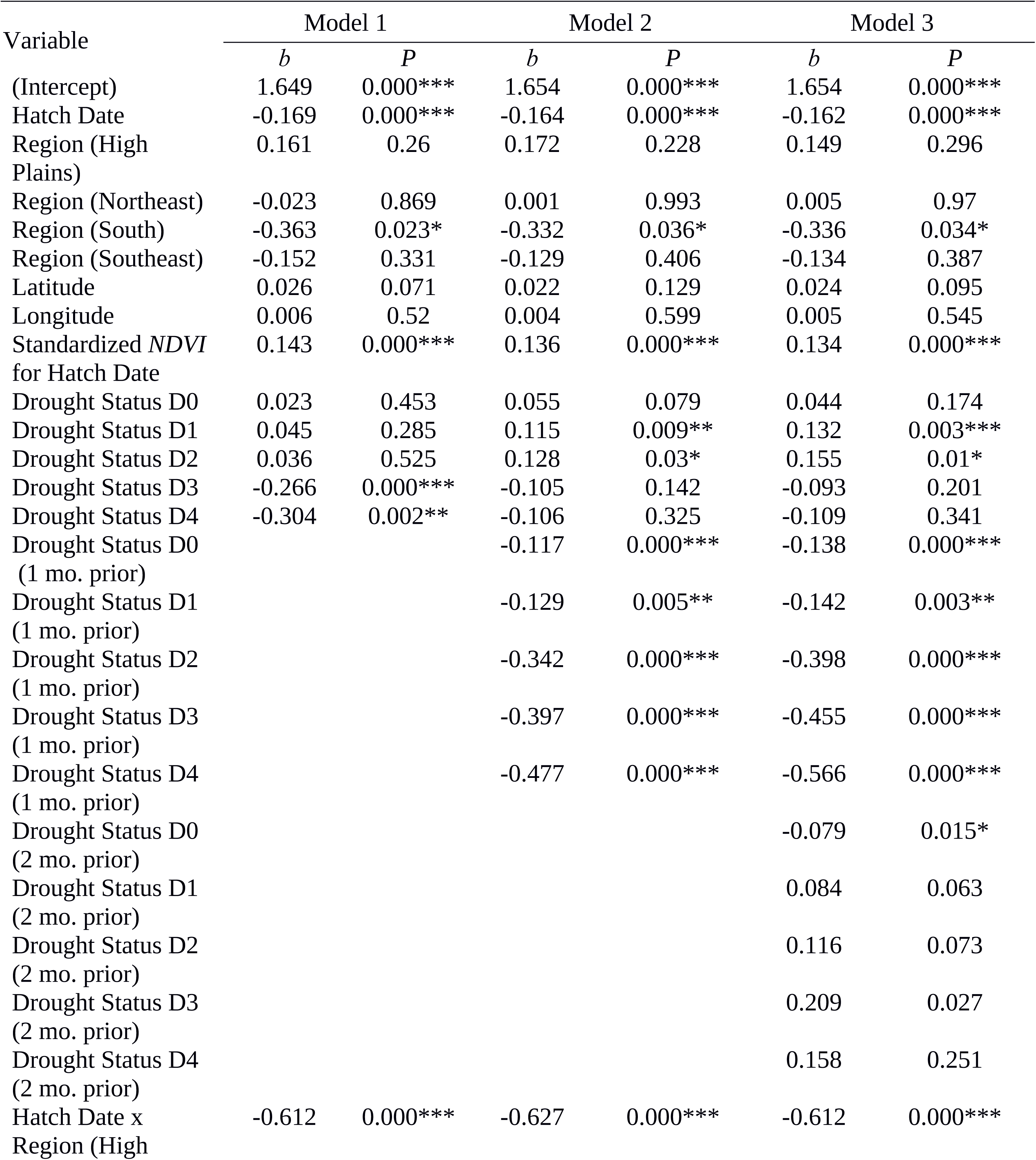
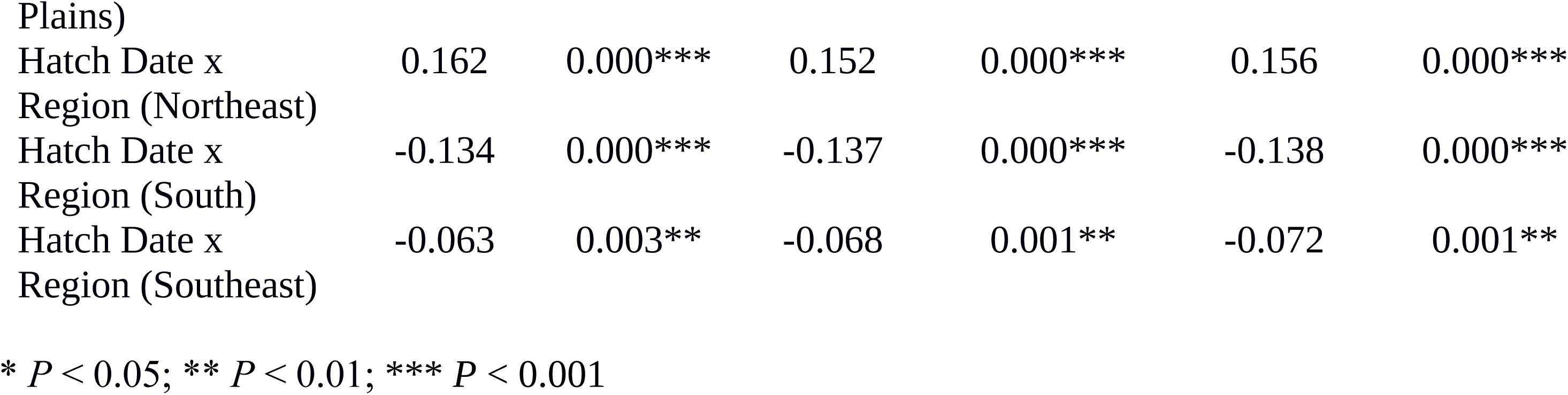
Parameter *β* estimates (*b*) and probabilities (*P*) for three generalized linear models used to evaluate drought effects on Eastern Bluebird hatch ratio. Explanatory variables are lay date, region, latitude, longitude, standardized *NDVI* for the hatch date, drought status during the hatch date, drought status one month prior to the hatch date, and the interaction of hatch date and region.

Table 4 summarizes the fitted regression coefficients (and *p*-values) for each of the three models, using hatching ratio as the response variable. By exponentiation of the coefficients reported in the table, we calculated how the odds of eggs hatching (*e^b^*) were affected by drought status, month of the year, NADM climatic region, standardized NDVI, and the interaction of month and region. For drought occurring within the month of hatching, only D1 and D2 had significant effects on hatching; both had positive and increasing effects. By exponentiation of the coefficients, the odds ratios of hatching (compared to status N) were 1.141 for D1 and 1.17 for D2. Drought at all levels within one month prior to hatching was highly significant and decreased hatching ratios as drought intensity increased. For drought status one month prior, the odds ratio of hatching declined steadily from 0.871 for D0 to 0.568 for D4. For drought occurring two months prior to hatching, only D0 had a significant effect with a negative effect on hatching; the odds ratio (0.924) was significantly lower compared to the reference of no drought. Hatch ratio was significantly affected by NDVI_std.,_ and the odds ratio of an egg hatching increased by a factor of 1.14 with each unit increase.

In addition to the drought-status effects, hatch date had significant effects on hatching ratios. For month, the odds of hatching decreased as the months progressed from March to August; for a one-unit increase in month, the odds ratio of an egg hatching decreased by a factor of 0.85. Only the South NADM climatic region was significant as an effect which suggested a decrease in hatching ratios and odds of hatching (0.71) compared to the reference region. Moreover, all interactions of hatch date and region were significant, but only for the Northeast region were the odds of hatching increased (1.17).

A seasonal decline in hatching was evident across all regions (Fig. 5). Following a peak in March and April, the mean number of eggs hatching decreased as the season progressed. There was an increase in hatching within the South region during the final month of the breeding season, but this was the case for only a very few observations.

#### Fledging rate

The modal number of fledging young was 4, followed by zero and 5 (Fig. 3). Again, the frequency distribution was bimodal, with more zero fledges than zero hatches. The average number fledged was 2.93. The number of fledged young declined from more than 3.0 fledges per nest to 2.4 fledges per nest with increasing drought status, especially for the two most extreme categories (Fig. 4).

As with clutch size and hatching ratio, we used a tiered approach, and summarize the results in Table 5. The results agreed closely with those for hatching ratio; LR tests favored models including current month and prior month drought statuses (*Χ*^2^ = 48.39, *df* = 5, *P* << 0.0001, *Χ*^2^ = 21.33, *df* = 5, *P* << 0.001, *Χ*^2^ = 69.8, *df* = 5, *P* < << 0.001, respectfully), and in favor of the more complex random-effect structure (*Χ*^2^ = 8550.7, *df* = 2, *P* << 0.0001).

**Table 5.**
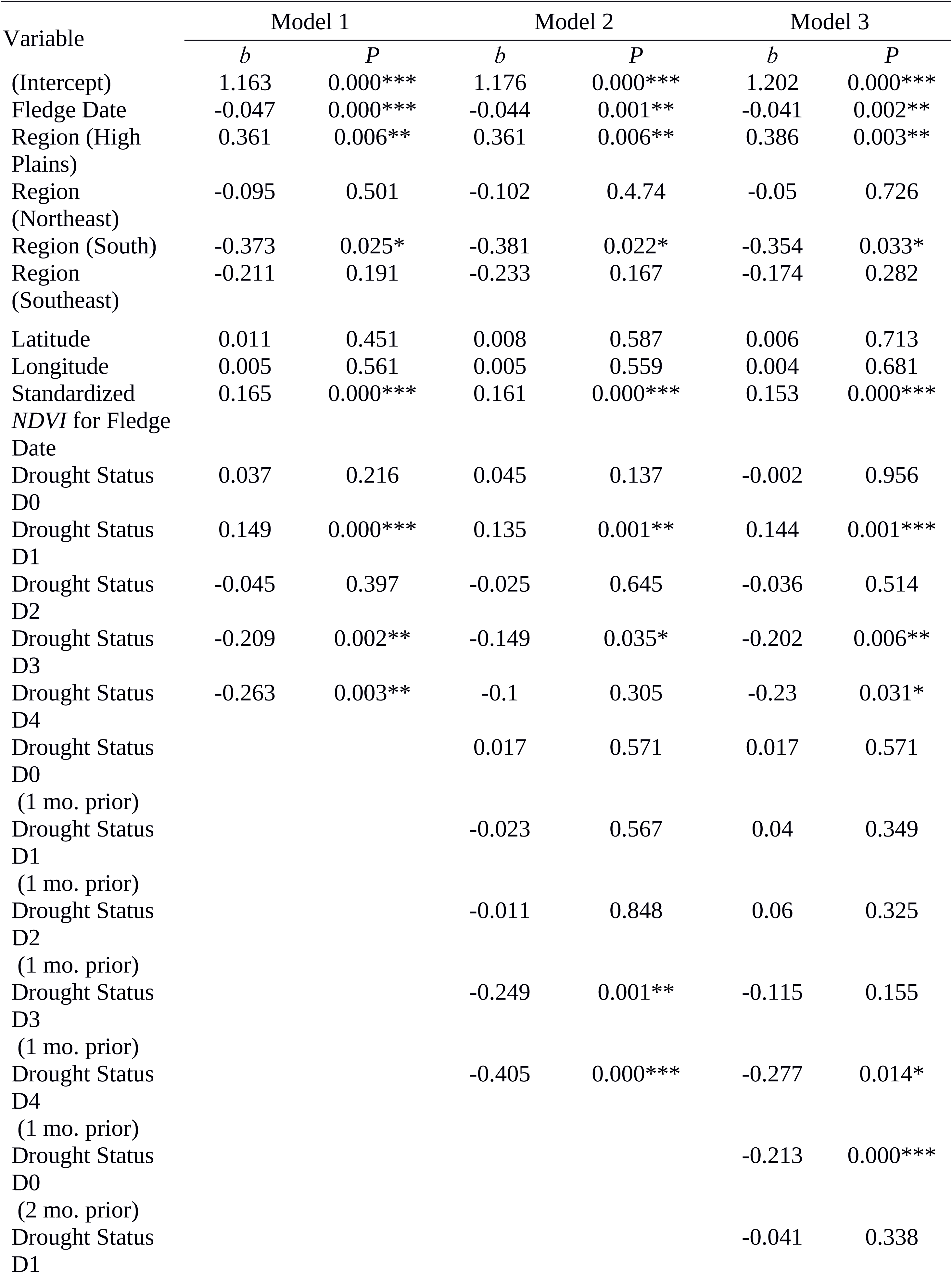
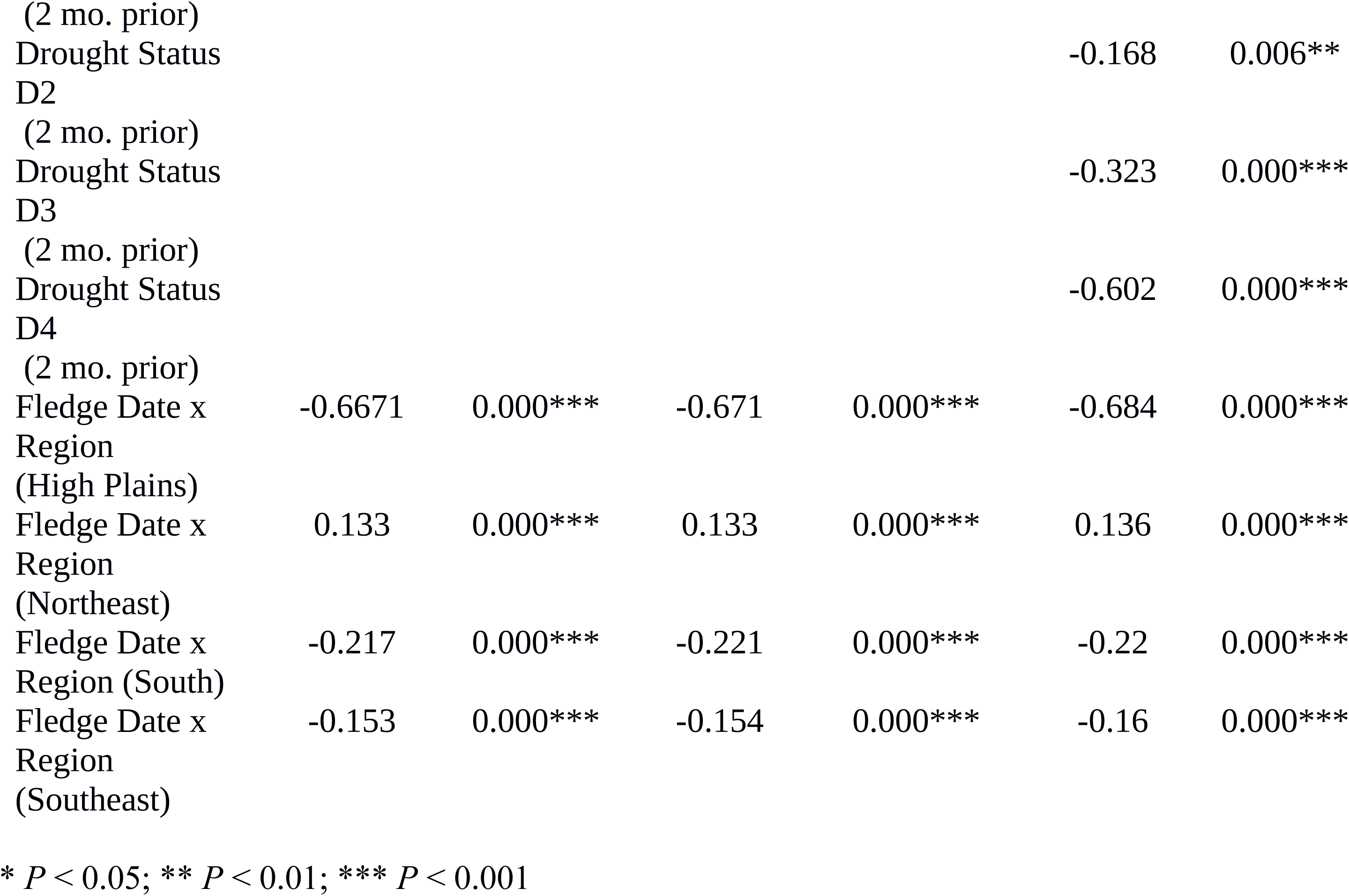
Parameter *β* estimates (*b*) and probabilities (*P*) for three generalized linear models used to evaluate drought effects on Eastern Bluebird fledging ratio. Explanatory variables are fledge date, region, latitude, longitude, standardized *NDVI* for the fledge date, drought status during the fledge date, drought status two months prior to the fledge date, and the interaction of fledge date and region.

We interpret the drought status results for Model 3, since our sequential hypothesis testing results were overwhelmingly in favor of including both the current and prior (one and two months) drought status variables in the model. The odds ratio of fledging increased by a factor of *e* ^-0. 144^ = 1.155 for current month drought status D1. For drought statuses D3 and D4 in the current month, we observed a decreasing effect on the odds of fledging within this model (0.817 and 0.795, respectfully). Drought status one month prior to fledging was only significant for the most extreme drought (D4) and decreased the odds of fledging (0.759). For drought statuses two months prior to fledging, all but D1 had significant negative effects on fledging; levels D0 and D2‒D4 decreased fledging odds (0.808, 0.845, 0.724, and 0.548, respectively). The odds ratio of fledging increased by a factor of *e* ^0.153^ = 1.165, with each unit increase in standardized NDVI.

Fledging rate generally decreased as the fledge date progressed through the breeding season. In addition, fledging rate increased significantly and was greater on the High Plains and decreased significantly and was less in the South, compared to the Midwest. The odds ratios of young fledging were 1.147 for the High Plains, 0.951 for the Northeast, 0.702 for the South, and 0.840 for the Southeast. Finally, there were significant interactions between fledge date and all of the regions, with decreased odds of fledging for all except the Northeast. As shown in Fig. 5, the mean number of young fledging generally declined as the season progressed. As with the mean number of eggs, the mean number of young fledging in the South region increased at the end of the breeding season (Fig. 5).

#### Spatial autocorrelation

To assess whether our model successfully accounted for spatial autocorrelation in the data, we tested for the presence of spatial autocorrelation in the model residuals using Moran’s I. Using the Moran.I function from the R library ape, we calculate the global Moran’s I for each of the years 2007 through 2013 (we left out 2006 due to low number of observations). The Moran’s I analysis indicated the presence of residual spatial autocorrelation at finer (within county) spatial scales. On a coarser between-county spatial scale, the analysis showed that the null hypothesis could not be rejected at level alpha = 0.05 (for all of the third-tier models), indicating that our models accounted for the spatial autocorrelation structure in the data at coarser scales. On average, there were 10 observations per county.

## Discussion

We found that drought affected Eastern Bluebird reproduction in both expected and unexpected ways. Drought had no effect on clutch size, regardless of occurrence within two months prior to and during clutch initiation. Severity of drought had profound effects on both hatching and fledging ratios, with the most severe levels (D3 and D4) resulting in the greatest decreases in both variables. Drought occurring one-month and two-months prior also had significant effects on hatching and fledging; moreover, the most severe drought levels amplified the decrease in hatching ratio typically seen as the breeding season progresses.

### Clutch size

Results of our initial exploratory analyses were supportive of increased clutch sizes with increasing latitude, which is typical of most passerine birds [26, 27]. Latitude was not significant when we included NADM climate regions in the final models, but we suspect this effect was diluted because the climate regions spanned multiple latitudes. As suggested by Crick et al. [37] and supported by Dhondt et al. [28] and Cooper et al. [20] migratory, multi-brooded species breeding in more northern latitudes, including Eastern Bluebirds, produce larger clutches earlier in the breeding season compared to non-migratory populations breeding in southern latitudes. Southern populations also tend to produce larger clutches mid-season. Our results also followed this trend, as we observed that clutch size decreased at a slower monthly rate in the South and Southeastern regions (relative to the Midwest). The increased mean clutch size observed for the South region at the end of the breeding season, and accompanying increases in mean number hatching and fledging, was most likely associated with the fewer number of observations reported from that region.

Eastern Bluebird clutch size was not affected by any level of drought or correlated with standardized NDVI. Likewise, clutch sizes of Vesper Sparrows (*Pooecetes gramineus*) were not affected during historically severe drought conditions within the breeding range of that species [3]. The non-sensitivity of clutch size to drought may be due to its unpredictable and variable nature and evolutionary adaptations that favor production of an optimal clutch size even during environmentally challenging years [38]. Factors other than environment, such as female fitness, predation of the incubating female or eggs, or inaccuracies in the counting of eggs by observers, may have also influenced clutch size data.

Eastern Bluebirds residing above 38^◦^ N generally undertake short distance migrations in winter, whereas more southern populations remain resident year round [30]. It was not possible for us to definitively determine which of the more northerly observation records were of migratory vs. non-migratory pairs. This restricted our examination of prior drought effects to only two months prior to clutch initiation, when pairs begin establishing territories and females commence building nests. As we documented, however, there was no drought effect on clutch size, though we suspect that migratory status would not contribute to the ability of a female to produce an optimal clutch.

### Hatching success

In this study, considerably more nests (55%) experienced a 100% hatch rate when there was no drought (N) than when there were the most severe of droughts (levels D3 and D4) (less than 40% hatch rate). Mild to severe drought (D0, D1, and D2) also appeared to influence hatch rate, but to a lesser degree.

The timing, and possibly severity, of drought within a single month likely varied on a week-by-week basis. The exact hatching date of any given clutch also varied within a single month given an expected average incubation period of 12 days. For example, clutches initiated earlier in a month should hatch during that month, while those initiated later would likely hatch in the following month. We elected not to include hatch date at a finer-scale than month because we felt the addition of additional variables would greatly decrease our ability to detect major trends.

In addition to the timing of clutch completion, it is possible that drought effects during the current month were muted when normal to mild drought conditions were in place during clutch formation in prior month. This may explain why current month drought statuses D1 and D2 had an unexpected positive influence on hatching ratios (relative to the hatching ratio at status N), while drought statuses D3 and D4 in the current month had no effect on hatching ratio. At least for nests under D1 drought during the expected month of hatching, nearly half of those (∼ 48%) had been under normal or abnormally dry conditions in the prior month. Drought status one month prior to hatching had a much more significant and negative effect as evidenced by the greatly lowered the odds of hatching under all drought statuses. Drought occurring two months prior had no or a very much reduced effect. This would be expected given that this time period would be well before clutch initiation.

In addition to the drought-status effects, there were highly significant interactions between month of hatching and region. The Northeast, the region least subject to drought statuses of D1 or greater, had a positive influence on hatch ratio when the interaction with hatch date was included, while those regions with the more frequent, more intense, and longer duration of drought (High Plains, South, and Southeast) had highly significant negative interactions with hatch month. Thus, drought one month prior to the month of hatching, hatch date, and interaction of hatch date and region had significant effects on the ratio of eggs that hatched.

The decreased hatching success documented in this study with increasing drought level occurring one-month prior to hatching may have resulted from factors associated with higher drought-associated temperatures during the pre-incubation period. We did not, however, include ambient temperature in our analyses. It is possible, although not necessarily so, that there were higher than normal temperatures occurring concurrently with drought conditions. For example, Karnieli et al. [39] demonstrated a negative relationship between NDVI and land surface temperature (LST) during the May to October period, which coincides with the latter portion of the breeding season. Changes in female incubation behavior, loss of egg viability, embryo mortality, or a combination are associated with higher ambient temperatures occurring before and during incubation [29, 40]. It has also been suggested that higher temperatures reduce egg viability via changes in embryo development, but not necessarily embryo mortality [29]. The egg-viability hypothesis suggests that egg viability varies with latitude, with clutch initiation date, and with the sequence at which eggs are laid [29]. In Swamp Sparrows (*Melospiza georgiana*) [41], Pearly-eyed Thrashers (*Margarops fuscatus*) [42], and Green-rumped Parrotlets (*Forpus passerine*) [43], prolonged exposure to high ambient temperature, as would occur with the first eggs laid in a clutch, decreased egg viability and reduced hatching success.

Month of hatching significantly influenced hatch ratios such that hatch rate declined as the breeding season progressed. The observed pattern of increased hatching failure later in the breeding season, and at lower latitudes, is not uncommon among many bird species [20, 44]. Indeed, it is well-documented that hatching failure of Eastern Bluebirds eggs follows this pattern [20, 28]. Our results suggest that this effect may be amplified during extreme and exceptional drought.

### Fledging success

Fledging success is influenced by many factors. Asynchronous hatching within a clutch, for example, results in a higher probability that the youngest nestling will die of starvation [45, 46]. Cooper et al. [29] found that female Eastern Bluebirds breeding at lower latitudes tended to initiate incubation before clutch completion, which would increase the likelihood of asynchronous hatching and loss of the youngest nestlings. Our observation of decreasing fledging success with increasing drought severity, especially when drought occurred 2 months prior to fledging, was possibly due to associated decreases in available food supply for both the nestlings and their parents. Smith [47] noted changes in insect densities and species composition along with a decline in numbers of insectivorous birds during drought. It is well established that the reproduction and abundance of herbivorous insects is highly sensitive to the availability and quality of plant foliage [48]. Although the effects of precipitation on insect reproduction and survival have not been extensively studied [49], Thacker et al. [50] found variable rainfall during a critical stage of aphid reproduction to have a negative effect on both foliage quality and aphid abundance. It is thus likely that the timing and severity of drought negatively impacted insect populations that are critical for nestling survival.

Increased incidences of predation by rat snakes (*Pantherophis* spp.), which are common predators of bird’s eggs and nestlings [51], may have also contributed to the decreased hatching and fledging success we documented during severe drought. Variation in small mammal abundance has been associated with increased nest predation rates by snakes and other predators, including small raptors [51, 52]. Small omnivorous mammals (e.g., squirrels) may also resort to predation of eggs and nestlings when their food resources are limited [53]. NestWatch observation records did not provide information about incidences of predation by snakes or other predators, so we can only speculate that this may have contributed to lower fledging rates during drought.

### The Power of Citizen Science Data

Reproductive data acquired over a 7-year period through NestWatch and from our own study site, and drought data acquired from the NADM, provided the opportunity to examine drought effects across the Eastern Bluebird North American breeding range. While smaller studies conducted solely by experienced researchers yield more accurate data and have value for exploring localized effects, the field time required for a broader study is prohibitive. For example, reproductive data collected from our study site expended over 840 field hours on average for each approximately 173-day field season. Data entry was also time intensive. The more than 24,000 NestWatch records we received were easily imported into a database and then scrubbed of questionable records using simple SQL queries. Consequently, our data set was robust and efficient in terms of sample size and time expenditure, respectively.

## Conclusion

Our results demonstrated distinct drought impacts on certain aspects of Eastern Bluebird reproduction. We found a clear association between occurrence of extreme or exceptional droughts during critical periods of the Eastern Bluebird nesting cycle and decreases in both hatching and fledging success. We found especially striking that one of our models suggested that drought status one and two months prior to expected hatching and fledging is more important in explaining the effect of drought than during the month in which hatching or fledging takes place.

Eastern Bluebirds are a species of least concern [29] and thus not likely to experience widespread changes in population densities due to drought alone. We suggest, however, that drought effects could be significantly more deleterious for species in decline, especially in light of climate change. Studies based on citizen science-generated data for other species combined with standardized climate date could provide additional support for this observation.

## Acknowledgements

We thank the volunteers who collected the nesting data for this study through The Cornell Laboratory of Ornithology’s Nest Box Network, The Birdhouse Network, NestWatch, the Smithsonian Migratory Bird Research Center’s Neighborhood NestWatch, and various state nest monitoring projects that have contributed their data. We also thank the many students who assisted with our own bluebird monitoring program at Berry College and acknowledge the contributions of the late Catherine Chamberlain Graham, who assisted with literature review during the early phases of this project.

## References

1. Stahle DW, Fye FK, Cook ER, Griffin RD. Tree-ring reconstructed megadroughts over North America since a.d. 1300. Climatic Change. 2007;83(1):133.

2. Seager R, Tzanova A, Nakamura J. Drought in the Southeastern United States: causes, variability over the last millennium, and the potential for future hydroclimate change. J Climate. 2009;22:5021–45.

3. George TL, Fowler AC, Knight RL, McEwen LC. Impacts of a severe drought on grassland birds in western North Dakota. Ecol Appl. 1992;2(3):275–84.

4. Mooij WM, Bennetts RE, Kitchens WM, DeAngelis DL. Exploring the effect of drought extent and interval on the Florida snail kite: interplay between spatial and temporal scales. Ecol Model. 2002;149(1):25–39.

5. Niemuth ND, Solberg JW, Shaffer TL. Influence of moisture on density and distribution of grassland birds in North Dakota. Condor. 2008;110(2):211–22.

6. Brenner FJ. The influence of drought on reproduction in a breeding population of red-winged blackbirds. Am Midl Nat. 1966;76(1):201–10.

7. Langin KM, Sillett TS, Yoon J, Sofaer HR, Morrison SA, Ghalambor CK. Reproductive consequences of an extreme drought for orange-crowned warblers on Santa Catalina and Santa Cruz Islands. In: Damiani CC, Garcelon DK editors. Proceedings of the 7th California Islands Symposium. Arcata: Institute for Wildlife Studies; 2009. pp. 293–300.

8. Albright TP, Pidgeon AM, Rittenhouse CD, Clayton MK, Flather CH, Culbert PD, et al. Effects of drought on avian community structure. Glob Chang Biol. 2010;16(8):2158–70.

9. Takekawa JE, Beissinger SR. Cyclic drought, dispersal, and the conservation of the snail kite in Florida: lessons in critical habitat. Conserv Biol. 1989;3(3):302–11.

10. Bolger DT, Patten MA, Bostock DC. Avian reproductive failure in response to an extreme climatic event. Oecologia. 2005;142(3):398–406.

11. Dai A. Drought under global warming: a review. Wiley Interdisciplinary Reviews: Climate Change. 2010;2(1):45–65.

12. Sarhadi A, Ausín MC, Wiper MP, Touma D, Diffenbaugh NS. Multidimensional risk in a nonstationary climate: Joint probability of increasingly severe warm and dry conditions. Science Advances. 2018;4(11):eaau3487.

13. Jetz W, Wilcove DS, Dobson AP. Projected Impacts of Climate and Land-Use Change on the Global Diversity of Birds. PLoS Biol. 2007;5(6):e157.

14. Langham GM, Schuetz JG, Distler T, Soykan CU, Wilsey C. Conservation status of North American birds in the face of future climate change. PLoS One. 2015;10(9):e0135350.

15. Lehikoinen A, Virkkala R. North by north-west: climate change and directions of density shifts in birds. Glob Chang Biol. 2016;22(3):1121–9.

16. Silvertown J. A new dawn for citizen science. Trends Ecol Evol. 2009;24(9):467–71.

17. Hochachka WM, Fink D, Hutchinson RA, Sheldon D, Wong W-K, Kelling S. Data-intensive science applied to broad-scale citizen science. Trends Ecol Evol. 2012;27(2):130–7.

18. McKinley DC, Miller-Rushing AJ, Ballard HL, Bonney R, Brown H, Cook-Patton SC, et al. Citizen science can improve conservation science, natural resource management, and environmental protection. Biol Conserv. 2017;208:15–28.

19. Sullivan BL, Phillips T, Dayer AA, Wood CL, Farnsworth A, Iliff MJ, et al. Using open access observational data for conservation action: A case study for birds. Biol Conserv. 2017;208:5–14.

20. Cooper CB, Hochachka WM, Phillips TB, Dhondt AA. Geographical and seasonal gradients in hatching failure in eastern bluebirds (*Sialia sialis*) reinforce clutch size trends. Ibis. 2006;148(2):221–30.

21. Dunn PO, Winkler DW. Climate change has affected the breeding date of tree swallows throughout North America. Proc Biol Sci. 1999;266(1437):2487-90.

22. Phillips T, Dickinson J. Tracking the nesting success of North America’s breeding birds through public participation in NestWatch. In: Rich TD, Arizmendi C, Demarest DW, Thompson, C, editors. Proceedings of the fourth international Partners in Flight conference. McAllen: Partners in Flight; 2009. pp. 633–40.

23. Cornell Lab of Ornithology. NestWatch. [cited 10 August 2017]. Available from: http://nestwatch.org

24. Pinkowski BC. Breeding adaptations in the eastern bluebird. Condor. 1977;79(3):289- 302.

25. Carleton RE, Pruett H. If you build them, they will come: nest boxes increase eastern bluebird recruitment within a site in northwest Georgia. Oriole. 2011;76(3-4):57–64.

26. Lack D. Ecological adaptations for breeding in birds. London: Methuen; 1968.

27. Ricklefs E. Geographical variation in clutch size among passerine birds: Ashmole’s hypothesis. Auk. 1980;97(1):38–49.

28. Dhondt AA, Kast TL, Allen PE. Geographical differences in seasonal clutch size variation in multi-brooded bird species. Ibis. 2002;144(4):646–51.

29. Cooper CB, Hochachka WM, Butcher G, Dhondt AA. Seasonal and latitudinal trends in clutch size: thermal constraints during laying and incubation. Ecology. 2005;86(8):2018–31.

30. Gowaty PA, Plissner JH. Eastern bluebird (Sialia sialis), version 2.0. In: Poole A, editor. The Birds of North America Online. Ithaca, NY: Cornell Lab of Ornithology; 2015.

31. Fair J, E. Paul, J. Jones, A.B. Clark, C. Davie, G. Kaiser, editors. Guidelines to the use of wild birds in research. 3rd ed. Washington: The Ornithological Council; 2010.

32. Kerr JT, Ostrovsky M. From space to species: ecological applications for remote sensing. Trends Ecol Evol. 2003;18(6):299–305.

33. Albright TP, Pidgeon AM, Rittenhouse CD, Clayton MK, Wardlow BD, Flather CH, et al. Combined effects of heat waves and droughts on avian communities across the conterminous United States. Ecosphere. 2010;1(5):art12.

34. Visser ME, Both C, Lambrechts MM. Global climate change leads to mistimed avian reproduction. Adv Ecol Res. 2004;35:89–110.

35. Fox J, Monette G. Generalized collinearity diagnostics. J Am Stat Assoc. 1992;87(417):178–83.

36. Nielsen-Gammon JW. Office of the State Climatologist Report: The 2011 Texas Drought. College Station: Texas A&M University; 2011.

37. Crick HQP. The impact of climate change on birds. Ibis.146(s1):48–56.

38. Boyce MS, Perrins CM. Optimizing great tit clutch size in a fluctuating environment. Ecology. 1987;68(1):142–53.

39. Karnieli A, Agam N, Pinker RT, Anderson M, Imhoff ML, Gutman GG, et al. Use of NDVI and land surface temperature for drought assessment: merits and limitations. J Climate. 2010;23:618–33.

40. Conway CJ, Martin TE. Effects of ambient temperature on avian incubation behavior. Behav Ecol. 2000;11(2):178–88.

41. Olsen BJ, Felch JM, Greenberg R, Walters JR. Causes of reduced clutch size in a tidal marsh endemic. Oecologia. 2008;158(3):421.

42. Beissinger SR, Cook MI, Arendt WJ. The shelf life of bird eggs: testing egg viability using a tropical climate gradient. Ecology. 2005;86(8):2164–75.

43. Stoleson SH, Beissinger SR. Egg viability as a constraint on hatching synchrony at high ambient temperatures. J Anim Ecol. 1999;68(5):951–62.

44. Koenig WD. Ecological and social factors affecting hatchability of eggs. Auk. 1982;99:526–36.

45. Lack D. The significance of clutch-size. Ibis. 1947;89(2):302–52.

46. Slagsvold T, Wiebe KL. Hatching asynchrony and early nestling mortality: the feeding constraint hypothesis. Anim Behav. 2007;73(4):691–700.

47. Smith KG. Drought-induced changes in avian community structure along a montane sere. Ecology. 1982;63(4):952–61.

48. Walther G-R, Post E, Convey P, Menzel A, Parmesan C, Beebee TJC, et al. Ecological responses to recent climate change. Nature. 2002;416(6879):389–95.

49. Bale JS, Masters GJ, Hodkinson ID, Awmack C, Bezemer TM, Brown VK, et al. Herbivory in global climate change research: direct effects of rising temperature on insect herbivores. Glob Change Biol. 2002;8(1):1–16.

50. Thacker JI, Thieme T, Dixon AFG. Forecasting of periodic fluctuations in annual abundance of the bean aphid: the role of density dependence and weather. J Appl Entomol. 1997;121(1-5):137–45.

51. Fitch HS. Natural history of the black rat snake (*Elaphe o. obsoleta*) in Kansas. Copeia. 1963;4:649–58.

52. Schmidt KA, Ostfeld RS. Songbird populations in fluctuating environments: predator responses to pulsed resources. Ecology. 2003;84(2):406–15.

53. Clotfelter ED, Pedersen AB, Cranford JA, Ram N, Snajdr EA, Nolan V, et al. Acorn mast drives long-term dynamics of rodent and songbird populations. Oecologia. 2007;154(3):493–503.

